# Glioma-BioDP: Database for Visualization of Molecular Profiles to Improve Prognosis of Brain Cancer

**DOI:** 10.1101/2022.06.09.495541

**Authors:** Shaoli Das, Xiang Deng, Harpreet Kaur, Evan Wilson, Kevin Camphausen, Uma Shankavaram

## Abstract

Cancer researchers often seek user-friendly interactive tools for validation, exploration, analysis, and visualization of molecular profiles in cancer patient samples. To aid researchers working on both low-and high-grade gliomas, we developed Glioma-BioDP, a web tool for exploration and visualization of RNA and protein expression profiles of interest in these tumor types. Glioma-BioDP is an extended version of our earlier published tool GBM-BioDP. In this version we have included expression data from both the low-and high-grade glioma patient samples from The Cancer Genome Atlas and enabled querying by mRNA, microRNA, and protein level expression data from Illumina HiSeq and RPPA platforms respectively. Glioma-BioDP enables users to explore the association of genes, proteins, and miRNA expression with molecular and/or histological subtypes of gliomas, surgical resection status and survival. The prognostic significance and visualization of the selected expression profiles can be explored using interactive utilities provided. This tool also enables potential validation and generation of new hypotheses of novel therapies impacting gliomas that aid in personalization of treatment for optimum outcomes.

**Availability:** Glioma-BioDP web tool with user manual is available from: https://glioma-biodp.nci.nih.gov Contact

## Introduction

Gliomas are the most common types of brain cancers originating in the glial cells. Initially the adult diffuse gliomas were classified according to the microscopic resemblance of the tumor cells with the normal glial cells [1]. In 2000, the World Health Organization (WHO) classified diffuse gliomas into the histological subtypes: astrocytomas, oligodendrogliomas, and oligoastrocytomas [2]. These were then graded for their degree of malignancy. Oligoastrocytomas and oligodendrogliomas are graded into grade II or III, while astrocytomas are graded into grades II, III and IV, the grade IV being known as glioblastomas [3]. Current classification of gliomas is based on genomic alterations in addition to the histopathological classification which are indicative of the aggressiveness of the tumor and patient prognosis [4]. Particularly, for lower grade gliomas (stage II, III), the mutation status of the isocitrate dehydrogenase 1 or 2 (*IDH1/2*) genes and codeletion in 1p19q is considered the key factors for molecular subtyping.

The availability of multi-omic cancer patient data from The Cancer Genome Atlas (TCGA) project has provided cancer researchers with unprecedented opportunities to explore and analyze molecular profiles in relation to patient survival, cancer stage, metastatic state, and other clinical factors [5]. However, downloading the bulk data and writing scripts for analysis and visualization of the data is a daunting task for most clinicians. Web tools like cBioPortal addresses these issues by incorporating interactive visualization and exploration of genetic profiles in different cancers [6]. However, cBioPortal has a generic interface for all tumor types. So, exploring the gene expression profiles in relation to the tumor-specific clinical parameters like molecular subtypes or specific driver alteration for a particular cancer of interest is not possible. To aid the cancer researcher working on gliomas we developed Glioma-BioDP, as an extended version of our previously published web tool GBM-BioDP [7]. Glioma-BioDP enables users with enhanced functions to query gene, protein, and miRNA expression profiles related to molecular subtypes, driver gene alteration status, histological subtypes as well as surgical resection status and patient survival.

## Materials and Methods

### Experimental and clinical data

In the current version of Glioma-BioDP, we collected RNA-seq (v2) and miRNA-seq data (level 3) from TCGA via Genomic Data Commons (GDC) data portal (https://portal.gdc.cancer.gov/). The tumor sample data used in this study is publicly available from TCGA, thus did not require approval from an ethics committee. We normalized the gene expression count data from RNA-seq and miRNA-seq platforms into log counts per million using the R package edgeR [8]. The normalized data was used for generating all the statistics we show for the expression distribution characteristics. The raw RPPA protein expression data were downloaded from (https://tcpaportal.org/tcpa/). We obtained both level 3 and level 4 data. All the protein expression data were processed and analyzed in a similar way.

### Database Construction

In the back end, we built a MySQL relational database. All the data presented by the Glioma-BioDP portal are retrieved from the in-house database that hosts information on clinical annotation, subtype information, gene, protein, and miRNA expression. The database also stored all the essential metadata, including the gene, miRNA, and sample information. The patient stratification showed in the portal for both TCGA-GBM and TCGA-LGG is based on the annotation reported by Verhaak and colleagues [9].

The Glioma-BioDP is a PHP based web application. The runtime high level architecture is 3-tiered, consistent with our previous released GBM-BioDP [7]. Processing is done in Python (http://www.python.org/) and visualization is developed using R (http://www.r-project.org/). The application is deployed on a secured Apache HTTP server (http://httpd.apache.org/) at the National Cancer Institute (NCI). The application will be maintained by periodic security scans for vulnerabilities and bugs.

## Results

### Modules

The Glioma-BioDP webtool has three modules: a) GBM, b) LGG, and c) GBM vs LGG as shown in Figure1a. Within each module, sub-modules exist to explore expression profiles and clinical details. Sub-module “Genes” include both mRNA and protein level expression from the TCGA Illumina HiSeq and RPPA platforms respectively. Sub-module “MIRNAs” contain expression data from TCGA Illumina HiSeq platform. The details of the modules are described below.

**Figure 1.**
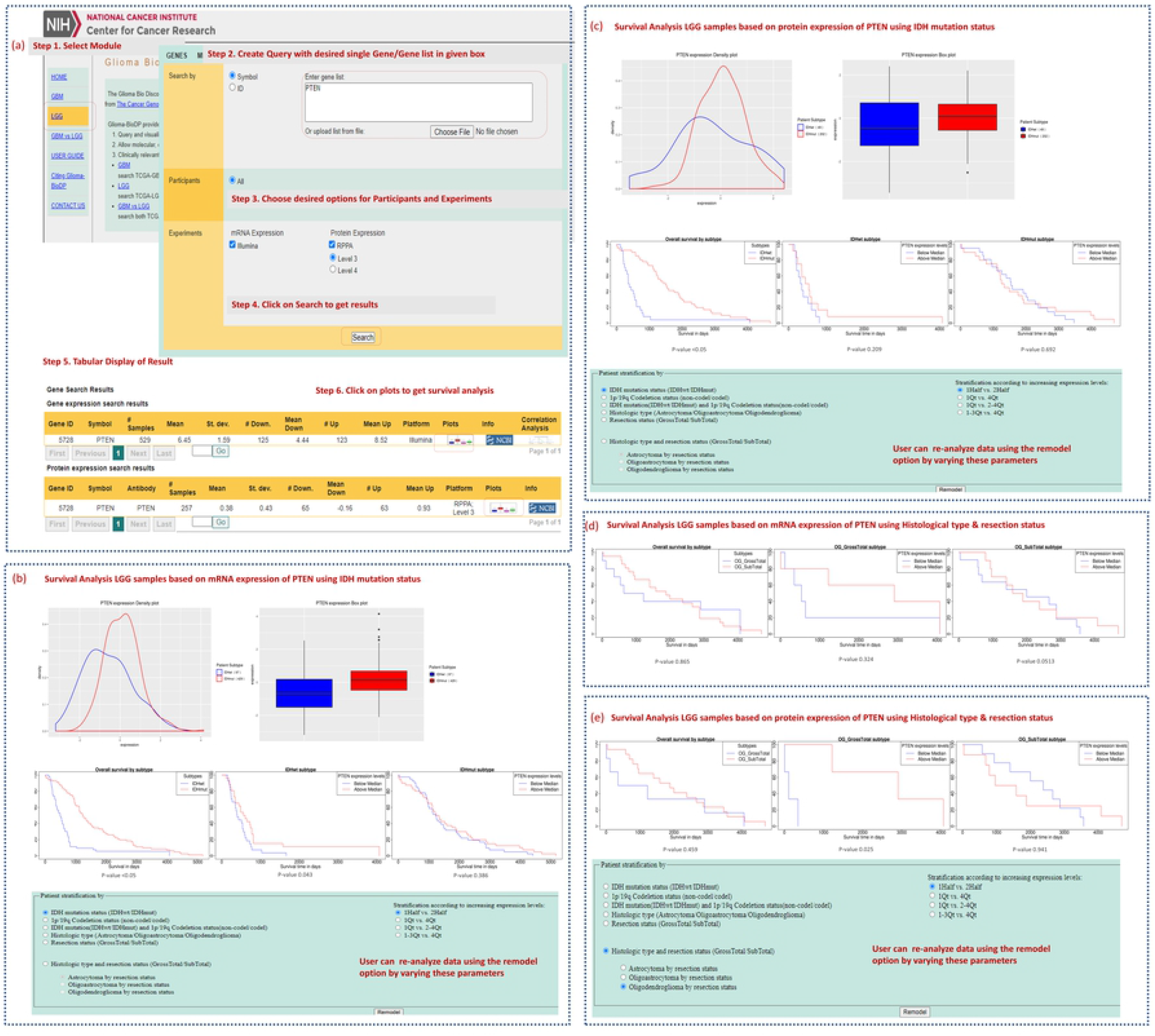
Workflow representing the steps involving the analysis of a gene (*PTEN*) to understand it’s prognostic implication in the LGG using the LGG module of Glioma-BioDP. (a) At step 1, the user can select either the LGG, GBM or LGG vs GBM module. At step 2 the user queries for gene (s) of interest followed by choosing appropriate participants and experiment types for mRNA expression and/or protein expression. At step 3, options are available for the specified gene with chosen participants and experiments. At step 4-5, results are displayed in the table. Next, at step 6, users can click on the plots icon to visualize the survival analysis results for the chosen gene. (b) Survival analysis results for *PTEN* mRNA and (c) protein expression respectively, in LGG based on IDH mutation status. (e) Survival analysis of *PTEN* mRNA and (f) protein expression respectively, in LGG based on histological type and resection status.

### Core Features

1) GBM module: Functionalities and options has been previously described in our publication [7].
2) LGG module: This module provides the visualization of expression profiles of LGG based on queried genes, proteins, or miRNAs. The visualization panel represents patient stratification with several options as follows:
  a) *IDH* wild type (*IDH*wt) vs. *IDH* mutation (*IDH*mut)
  b) 1p/19q codeletion status
  c) *IDH* and 1p/19q codeletion status
  d) histological subtypes including Astrocytoma, Oligoastrocytoma, Oligodendroglioma
  e) surgical resection status of gross total or sub-total.
  f) Histology and surgical resection for each of the LGG subtypes.
  For all the options mentioned above, Kaplan-Meier survival plots of expression profiles can be stratified by greater vs. less than median, and four expression quartiles (1st vs 4th, 1-2 vs 3-4, 1st vs 2-4, 1-3 vs 4th). The visualization of gene (or protein), or miRNA expression profiles between any of the above-mentioned stratifications is shown with a density plot and a box plot. The p-value for the difference of expression levels between these stratifications are calculated using t-tests. Prognostic significance of the gene or miRNA expression in any of the above-mentioned stratification is visualized with Kaplan-Meier survival curves stratified by the options as described above.
3) GBM vs LGG module: This module allows to explore prognostic significance of gene or miRNA expression in GBM vs. LGG. Visualization is shown in a side-by-side comparison with Kaplan-Meier survival plots. For the user selected genes or miRNA queries, the patient samples can be stratified by as the options described above.

### Workflow and Applications of Glioma-BioDP

Glioma-BioDP facilitates the user to assess the expression pattern and prognostic potential of desired gene, protein, or miRNA in specific brain tumor, i.e. GBM or LGG or both. Here, we describe the brief workflow of analyses using a clinically relevant gene *PTEN* as an example. The tumor suppressor gene *PTEN* plays important roles in the regulation of cell proliferation, apoptosis, and DNA damage repair [13]. Treatment of *PTEN*-deficient tumors with *PI3K* pathway inhibitors are being investigated for some cancer types [14]. The loss of *PTEN* expression has been indicated to be an early event in glioma, with mutations occurring in between 5% and 40% of glioma cases.

To get comprehensive understanding about the role of a selected gene (e.g. *PTEN*) in LGG subtypes from diverse perspective, we have integrated different sub-modules such as mRNA and protein expression based query of specific gene along with molecular and histological subtypes-based stratification of LGG tumors. Complete workflow for the analysis of *PTEN* in molecular and histological subtypes of LGG using LGG module is represented in the Figure 1a. Here, we have selected mRNA and protein expression-based query under the LGG module for the analysis of *PTEN* (steps 1-4). Glioma-BioDP allows the user to select any of desired options provided on the platform. Subsequently, the user will be directed to the tabular display of the gene and protein expression pattern in LGG samples. By clicking on the plots the user will be directed to the graphical interface (see steps 5-6 of Figure 1a). For demonstration purpose, we have used the option for *IDH* mutation. First, we observed *PTEN* mRNA expression in *IDH*mut is significantly (p-value <0.05) higher than that of *IDH*wt samples (boxplot panel of Figure 1b). Figure 1b is also showing three KM survival plots for: all LGG samples, *IDH*wt, and *IDH*mut samples respectively. The user can further select stratification options of survival risk groups in total LGG samples as well as LGG samples from IDH mutation subtype. Like the mRNA expression-based query, protein expression-based query can be performed to see differences in survival between different molecular subtypes of LGG (Figure 1c). Importantly, Glioma-BioDP allows the user to rebuild their survival models employing different parameters based on molecular features: *IDH* mutation and 1p/19q codeletion status, histological subtypes, surgical resection status, varying quartile ranges, etc. The resulting KM plots from the stratification of samples using histological type and resection status for mRNA and protein expression-based queries are shown in Figure 1d and 1e respectively.

### Case Studies

The Glioma-BioDP tool’s functionality and potential clinical relevance is demonstrated in LGG, GBM and GBM vs. LGG modules using the analyses of the following genes: *PTEN, NES, TERT, MGMT*,EGFR,. *NES* is a gene that codes for nestin, an intermediate filament found in vascular endothelial cells that is upregulated in tumors to allow for increased angiogenesis at the tumor site [11]. *TERT* codes for telomerase reverse transcriptase, a protein key to the maintenance of telomeres and one whose expression is upregulated in a subset of gliomas through promoter mutation or by other means to facilitate tumor progression [12]. *MGMT* promoter methylation and subsequent *MGMT* gene inactivation is common in malignant gliomas. Epigenetic *MGMT* promoter methylation has been shown to be associated with better clinical outcomes for patients treated with temozolomide (TMZ) and radiotherapy due to a decrease in tumor DNA repair [10]. This observation has great clinical relevance in terms of patient selection for chemo and radiotherapy. *EGFR* amplification and mutation are a signature genetic abnormality in GBM [15] and may be explored as a therapeutic target. Using these examples, we described the prognostic significance of each of these genes in LGG and GBM.

#### Prognostic value of the expression of *PTEN* in LGG subtypes

Expanding on the *PTEN* gene search in LGG module, the potential clinical relevance of overexpression of *PTEN* mRNA level was shown with Kaplan-Meier analysis (samples stratified by >median or <median of *PTEN* expression). In Figure 2a it is shown to be associated with better survival in *IDH*wt (p-value = 0.043) but not in *IDH*mut (Figure 2b). Also, querying *PTEN* expression in 1p19q co-deleted vs non-co-deleted LGGs showed that *PTEN* overexpression is associated with better survival in 1p19q non-co-deleted LGGs (Figure 2c, p-value = 0.004) but not in 1p19q co-deleted LGGs (Figure 2d). Further, querying for histological subtypes of LGG, it was seen that *PTEN* overexpression was associated with better survival in all histological subtypes: oligodendroglioma, astrocytoma, and oligoastrocytoma (Figure 2e-g, p-value = 0.03, 0.024 and 0.014 respectively). There are investigations underway for finding treatment strategies for targeting *PTEN*-deficient cancers. Association of *PTEN* overexpression with improved survival in glioma subtypes with poor prognosis (*IDH*wt and 1p19q non-co-deleted) may provide rationale for investigating the effects of these therapies in these glioma subtypes.

**Figure 2.**
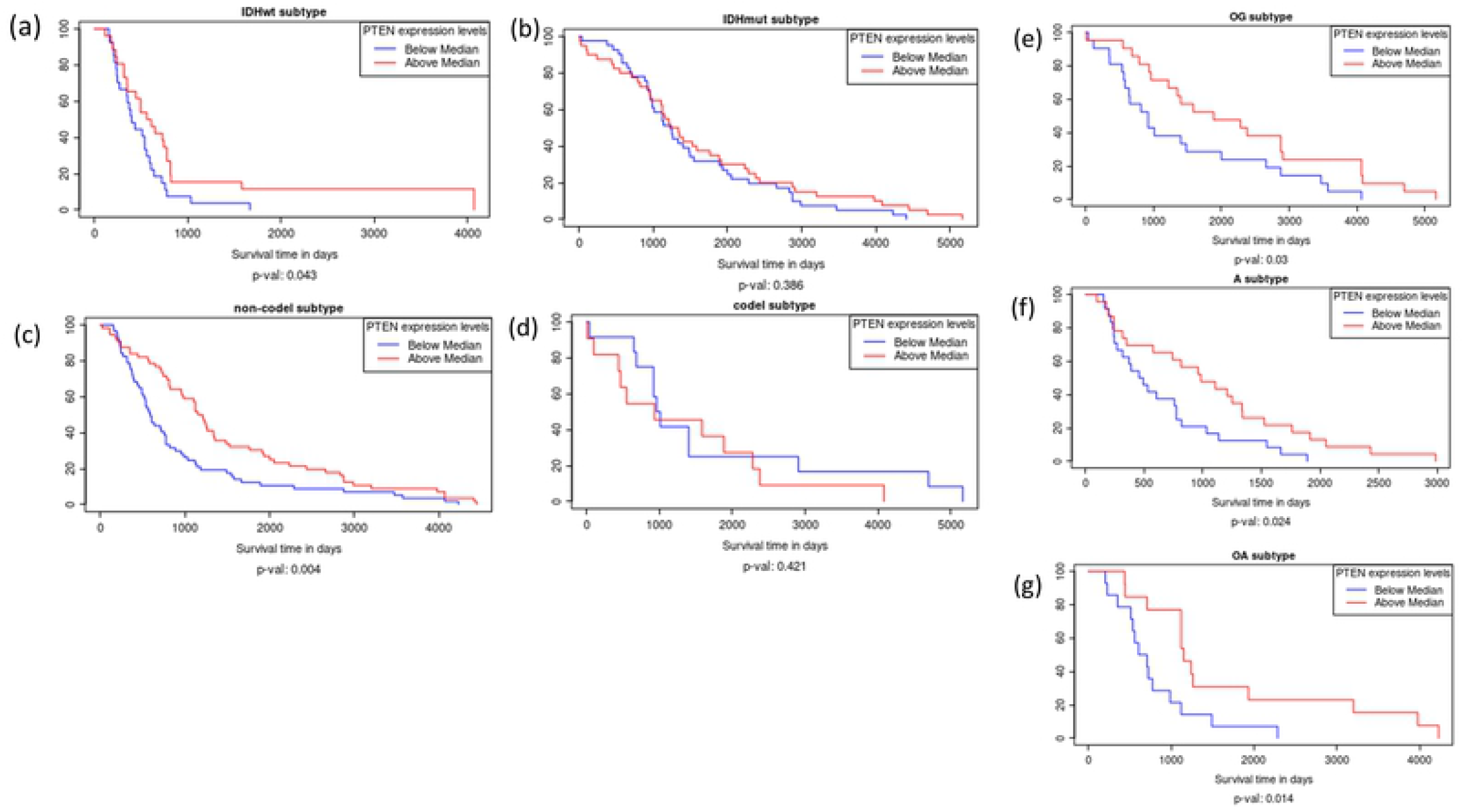
Kaplan Meier plots showing the clinical relevance of IDH and 1p19q status in LGG subtypes. (a,b) *PTEN* expression related to survival in *IDH*wt compared to *IDH*mut LGG. (c,d) *PTEN* expression related to survival in 1p19q non-co-deleted compared to 1p19q co-deleted LGG. (e) *PTEN* expression related to survival in oligodendroglioma. (f) *PTEN* expression related to survival in astrocytoma. (g) PTEN expression related to survival in oligoastrocytoma.

#### Prognostic value of the expression of genes *NES, TERT* and *MGMT* in LGG vs GBM

The webtool can showcase genes that are manipulated either in LGG, GBM or both tumor types. *NES* is an example of a gene that elucidates the Glioma-BioDP webtool’s ability to identify genes that have a significant effect on prognosis in LGG but not GBM. Comparing the 1^st^ vs. 4^th^ quartile of expression, elevated *NES* expression levels are shown to be associated with decreased survival times in LGG (Figure 3a, p-value: 0.003) but not GBM (Figure 3b, p-value: 0.998). *TERT* is an example of the genes that have prognostic significance in both LGG and GBM. Comparing the 1^st^ vs. 4^th^ quartile of expression, *TERT* expression levels are shown to have a similar effect on patient survival times in both LGG (Figure 3c-d, p-value: 0.043) and GBM (p-value: 0.036).

**Figure 3.**
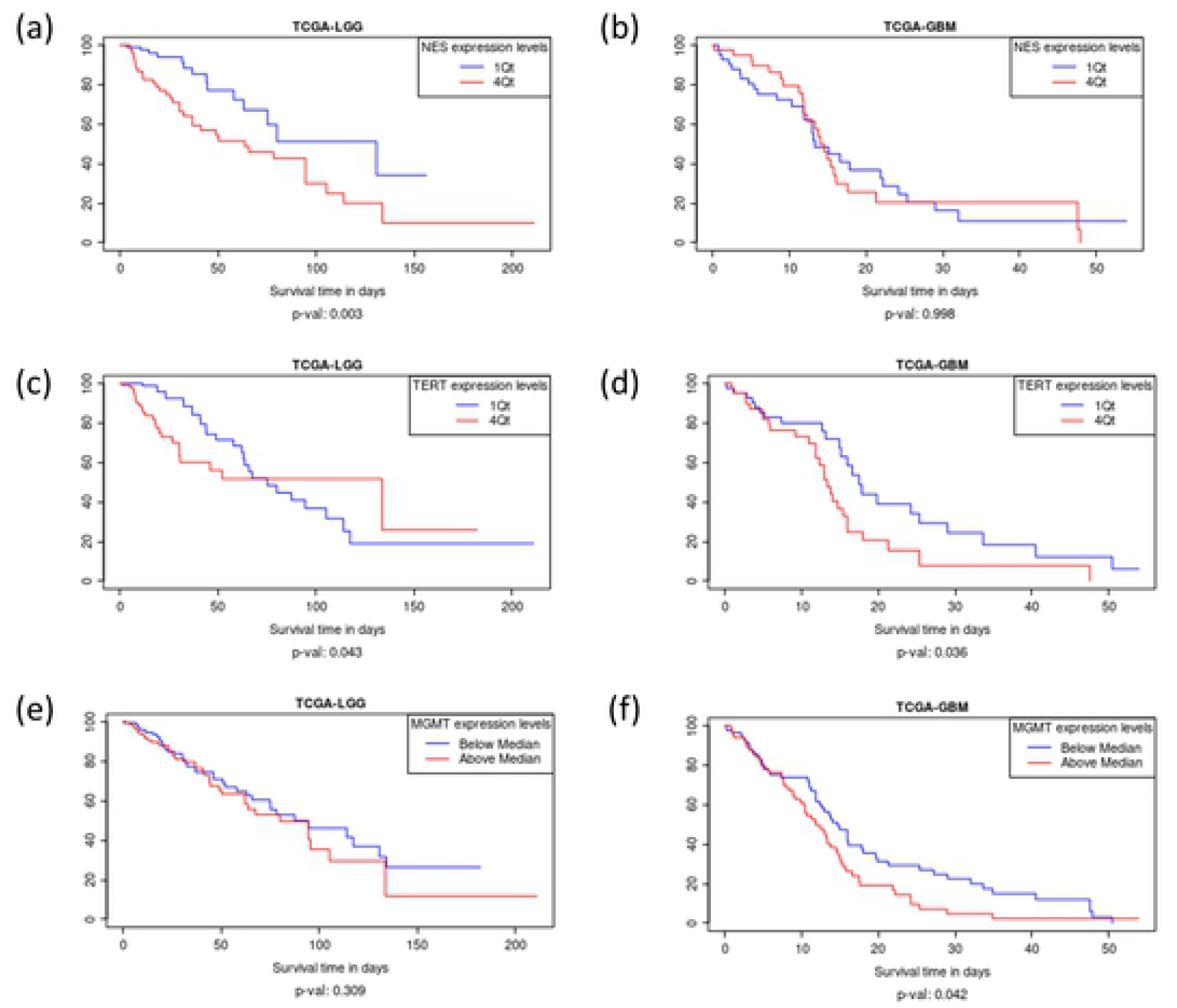
Kaplan Meier plots showing the comparison between LGG vs. GBM. Differences in protein expression changes in *NES* (a,b) and TERT (c,d) using the 1^st^ and 4^th^ quartiles option of survival plot. (e,f) showing the change in MGMT protein expression when selecting the median option for survival plot.

*MGMT* shows the webtool’s ability to highlight genes that show profound impact and significance in GBM but not LGG. The beneficial effect of *MGMT* promoter methylation and gene inactivation in GBM is corroborated by Glioma-BioDP. Stratifying *MGMT* expression by median, decreased *MGMT* protein expression not significantly associated with survival in LGG (Figure 3e, p-value: 0.309) but it is associated with longer survival times in GBM (Figure 3f, p-value: 0.026).

#### Prognostic value of the expression of *EGFR* in molecular subtypes of GBM and LGG

To explore the prognostic effects of *EGFR* expression in molecular subtypes of GBM and LGG, we queried Glioma-BioDP for each of these modules. When we queried for *EGFR* expression in GBM module, it could be observed that the mRNA and protein expression of *EGFR* is significantly higher in the classical subtype of GBM compared to all other subtypes (Figure 4a shows mRNA expression boxplot). Though patient stratification by *EGFR* expression within each molecular subtype do not show significant association with survival, in the proneural subtype strong trend is seen for association of better survival with overexpression of *EGFR* (Figure 4b, Kaplan-Meier analysis, p-value = 0.098, sample stratification by 1st quartile vs 2-4 quartiles). Patient survival in the other molecular subtypes of GBM did not show any association with *EGFR* expression level (Figure 4c-e). On the other hand, in LGG subtypes stratified by the presence of *IDH* mutation (*IDH*wt vs *IDH*mut), it could be seen that *EGFR* protein expression is significantly higher in *IDH*wt samples compared to *IDH*mut samples, as visualized using box plot (Figure 4f). From the Kaplan-Meier survival plots, it could be seen that overexpression of *EGFR* protein level is associated with better survival in *IDH*wt (Figure 4g), but not in the *IDH*mut subtype (Figure 4h) of LGG.

**Figure 4.**
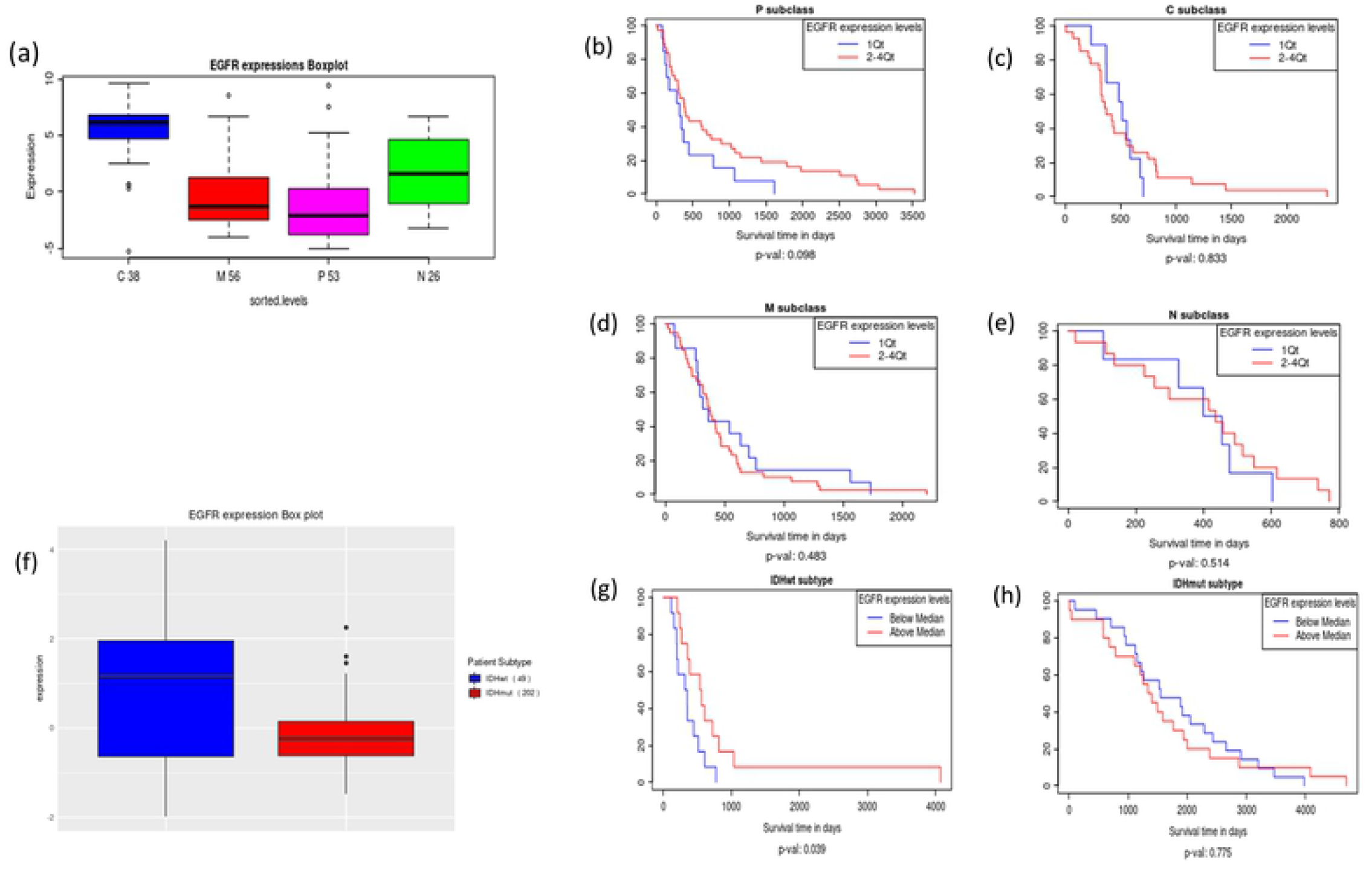
Comparison of EGFR changes GBM and LGG subtypes. (a) Boxplots of *EGFR* mRNA expression in GBM molecular subtypes. (b-e) *EGFR* mRNA expression related to survival in GBM molecular subtypes: (b) pro-neural (c) classical (d) mesenchymal, and (e) neural. (f) Boxplot view of *EGFR* protein expression in LGG subtype *IDH*wt compared to *IDH*mut. (g) *EGFR* protein expression related to survival in *IDH*wt subtype of LGG. (h) *EGFR* protein expression related to survival in *IDH*mut subtype of LGG.

Thus, the functionality of Glioma-BioDP is evident through the juxtaposition of these genes whose effect on patient outcomes is widely different yet similarly potent in either GBM or LGG or both settings. Also, querying Glioma-BioDP enables users to explore the mRNA and protein level profiles for their genes of interest in context of the molecular and histological subtypes.

### Summary and Future Direction

In the age of big data in cancer genomics, there is an opportunity for cancer researchers to use and explore the patient genomic data from large tumor cohorts such as TCGA, to improve their understanding of genomic correlates to patient prognosis. However, there is a need for availability of the data in easy to explore format and intuitive visualization that would enable the cancer researchers to make use of that enormous data. Glioma-BioDP as a user-friendly web tool offers intuitive visualization and query of gene and miRNA expression data in gliomas in context of specific histological and molecular subtypes of these tumors. In addition to our previously published tool GBM-BioDP, the new tool Glioma-BioDP enables exploration of the prognostic significance of transcriptomic and proteomic features from low grade to high grade gliomas in subtype-specific manner.

In comparison to a previously published tool GliomaDB [16], our tool Glioma-BioDP offers more intuitive and useful visualizations by enabling the users to look at gene or miRNA expressions between different histological, as well as molecular subtypes of gliomas. As described in our case studies, with Glioma-BioDP users get useful information on the survival of the glioma patients depending on the queried gene expression in context of the histological and/or molecular subtypes of gliomas. Inclusion of this information is a critical feature of Glioma-BioDP as the subtypes are linked to varied degree of patient prognosis in glioma, and the expressions of different genes may have different implications in prognosis depending on the glioma subtype. An example of this functionality is described by the association of *PTEN* overexpression with better survival in *IDH*wt and 1p19q non-co-deleted LGGs. Another example is that *EGFR* gene and protein level expressions are associated with prognosis in the glioblastomas (grade IV glioma), and even in the lower grade gliomas (grade II-III), *EGFR* protein level expression is associated better prognosis with the *IDH*wt molecular subtype which is the high-risk subtype compared to *IDH*mut.

In future versions of this tool, we are looking forward to integrating data from more omic platforms like mutation, copy number and DNA methylation that would expand the usability of this tool.

## Acknowledgements

This work was funded by the Intramural Research Program of the National Institutes of Health, National Cancer Institute.

## Ethics Statement

The tumor sample data used in this study is publicly available from TCGA. We used TCGA level 3 data that is devoid of the patient sensitive information, thus did not require approval from an ethics committee.

## References

1. Louis DN, Holland EC, Cairncross JG. Glioma classification: a molecular reappraisal. Am J Pathol 2001;159(3):779–86 doi: 10.1016/S0002-9440(10)61750-6[published Online First: Epub Date].

2. Gonzales M. The 2000 World Health Organization classification of tumours of the nervous system. J Clin Neurosci 2001;8(1):1–3 doi: 10.1054/jocn.2000.0829[published Online First: Epub Date].

3. Louis DN, Ohgaki H, Wiestler OD, et al. The 2007 WHO classification of tumours of the central nervous system. Acta Neuropathol 2007;114(2):97–109 doi: 10.1007/s00401-007-0243-4[published Online First: Epub Date].

4. Louis DN, Perry A, Reifenberger G, et al. The 2016 World Health Organization Classification of Tumors of the Central Nervous System: a summary. Acta Neuropathol 2016;131(6):803–20 doi: 10.1007/s00401-016-1545-1[published Online First: Epub Date].

5. Cancer Genome Atlas Research N, Weinstein JN, Collisson EA, et al. The Cancer Genome Atlas Pan-Cancer analysis project. Nat Genet 2013;45(10):1113–20

6. Cerami E, Gao J, Dogrusoz U, et al. The cBio cancer genomics portal: an open platform for exploring multidimensional cancer genomics data. Cancer Discov 2012;2(5):401–4

7. Celiku O, Johnson S, Zhao S, Camphausen K, Shankavaram U. Visualizing molecular profiles of glioblastoma with GBM-BioDP. PLoS One 2014;9(7):e101239 doi: 10.1371/journal.pone.0101239[published Online First: Epub Date].

8. Robinson MD, McCarthy DJ, Smyth GK. edgeR: a Bioconductor package for differential expression analysis of digital gene expression data. Bioinformatics 2010;26(1):139–40 doi: 10.1093/bioinformatics/btp616[published Online First: Epub Date].

9. Ceccarelli M, Barthel FP, Malta TM, et al. Molecular Profiling Reveals Biologically Discrete Subsets and Pathways of Progression in Diffuse Glioma. Cell 2016;164(3):550–63 doi: 10.1016/j.cell.2015.12.028[published Online First: Epub Date].

10. Hegi ME, Diserens AC, Gorlia T, et al. MGMT gene silencing and benefit from temozolomide in glioblastoma. N Engl J Med 2005;352(10):997–1003 doi: 10.1056/NEJMoa043331[published Online First: Epub Date].

11. Maderna E, Salmaggi A, Calatozzolo C, Limido L, Pollo B. Nestin, PDGFRbeta, CXCL12 and VEGF in glioma patients: different profiles of (pro-angiogenic) molecule expression are related with tumor grade and may provide prognostic information. Cancer Biol Ther 2007;6(7):1018–24 doi: 10.4161/cbt.6.7.4362[published Online First: Epub Date].

12. Brennan CW, Verhaak RG, McKenna A, et al. The somatic genomic landscape of glioblastoma. Cell 2013;155(2):462–77 doi: 10.1016/j.cell.2013.09.034[published Online First: Epub Date].

13. Smith JS, Tachibana I, Passe SM, et al. PTEN mutation, EGFR amplification, and outcome in patients with anaplastic astrocytoma and glioblastoma multiforme. J Natl Cancer Inst 2001;93(16):1246–56 doi: 10.1093/jnci/93.16.1246[published Online First: Epub Date].

14. Dillon LM, Miller TW. Therapeutic targeting of cancers with loss of PTEN function. Curr Drug Targets 2014;15(1):65–79 doi: 10.2174/1389450114666140106100909[published Online First: Epub Date].

15. Taylor TE, Furnari FB, Cavenee WK. Targeting EGFR for treatment of glioblastoma: molecular basis to overcome resistance. Curr Cancer Drug Targets 2012;12(3):197–209 doi: 10.2174/156800912799277557[published Online First: Epub Date].

16. Yang Y, Sui Y, Xie B, Qu H, Fang X. GliomaDB: A Web Server for Integrating Glioma Omics Data and Interactive Analysis. Genomics Proteomics Bioinformatics 2019;17(4):465–71 doi: 10.1016/j.gpb.2018.03.008[published Online First: Epub Date].

